# Gradient-k: Improving the Performance of K-Means Using the Density Gradient

**DOI:** 10.1101/2022.03.30.486343

**Authors:** Andrei Ciuparu, Raul C. Mureșan

**Affiliations:** Transylvanian Institute of Neuroscience, Department of Experimental and Theoretical Neuroscience, Pta. Timotei Cipariu 9/20, 400191 Cluj-Napoca, Romania; Technical University of Cluj-Napoca, Basis of Electronics Department, Str. G. Baritiu 26-28, 400027 Cluj-Napoca, Romania

**Keywords:** Clustering, k-means, density gradient, spike sorting

## Abstract

We introduce Gradient-k, an upgrade of the k-means algorithm that improves clustering accuracy and reduces the number of iterations required for convergence. This is achieved by correcting the distance used in the k-means algorithm by a factor based on the angle between the density gradient and the direction to the cluster center. We show that this correction allows the algorithm to go beyond the traditional tessellation obtained by k-means, enabling the creation of nonlinear separation boundaries. The correction also provides the ability to distinguish clusters having non-normal shapes and of varying densities. Furthermore, Gradient-k has a lower computational complexity for datasets with many samples. We compare the performance of Gradient-k with that of k-means and of DBSCAN on several benchmark datasets, as well as on a more realistic dataset in the context of neural spike sorting.

## Introduction

One of the hardest problems in machine learning pertains to unsupervised learning, which frequently involves the automatic identification of groups of samples (also called clusters or classes) in a dataset without having *a priori* knowledge about the number, shape, size, or location of these groups in space (1). Clustering algorithms must rely on statistical properties of the data, to make decisions on which samples are similar and group them together. One of the most popular clustering techniques is k- means (2), which has a relatively simple computational implementation and scales easily to multiple dimensions.

K-means falls into the class of distance-based algorithms, whereby data points are aggregated based on some distance measure. These algorithms suffer from the curse of dimensionality (3,4):as the dimensionality of the problem increases, distance begins to lose meaning. If the data is in a very high dimensional space, distances between samples increase and it is more difficult to make decisions based only upon Euclidian distance, for example. In its original form, k-means only uses the Euclidian distance between samples (datapoints) and cluster centers to decide whether the point belongs to the cluster or not, averages the location of all points assigned to a cluster to determine the new cluster centre location, and repeats this process iteratively until all cluster centers are stable. The stopping condition for k-means is generally phrased in terms of minimum cluster centre movement. The algorithm is computationally simple, easy to implement, and scales well to large datasets. However, it suffers from at least two major issues, as discussed below: k-means is sensitive to the initial conditions and it has issues related to the distribution of point density in data space.

Indeed, k-means is a stochastic clustering technique, as the solution may depend on the initial conditions (cluster centers). There are several algorithms for choosing the initial cluster centers, but the most widely used is the K++ initialization, first described in 2007 by David Arthur and Sergei Vassilvitskii (5). This algorithm attempts to spread out the initial clusters evenly across the data space, leading to an increased convergence speed and accuracy.

Furthermore, the main assumption of distance-based algorithms, like k-means, is that clusters are distributed in space according to a normal density distribution (6). This boils down to the idea that clusters are denser closer to their centre and less dense towards their exterior. Distance-based algorithms lose power when clusters have non-normal shapes, or if clusters are of varying sizes and varying densities. The same algorithms also create linear separation boundaries between neighboring clusters, failing to identify non-linear splits (7). If the cluster shapes are non-Gaussian, and/or the clusters are placed in close proximity or have part of their points surrounded by other clusters, then these algorithms tend to mislabel points because of tessellation (pairs of clusters are separated according to linear splits) (8,9).

Another class of clustering algorithms relies on the density of points in space (10), whereby points are grouped iteratively according to some local density criterion (11). These avoid the tessellation problem by being able to create non-linear separation boundaries and effectively reducing the Gaussian assumption to a thresholding operation. One of the weaknesses of density-based algorithms is that identifying clusters with different densities requires different density thresholds and, as such, they perform poorly when cluster densities are unbalanced.

Here, we present the Gradient-k algorithm, an improvement of the k-means implementation that uses the gradient of the point density function to improve accuracy and convergence speed. Our goal was to design an improved version of k-means that would solve some of its main issues by using more information present in the data. Inspired by density-based techniques, the new algorithm uses auxiliary information about how the data is distributed in space, enabling it to detect clusters regardless of their density, shape, and size. This is achieved by computing the gradient of an approximate density function in order to correct the distance metric used by k-means, effectively informing the distance metric about the distribution of the datapoints.

## Methods

### The Gradient-k Algorithm

While the Gradient-k algorithm could be generalized to N dimensions, here we only discuss results on two-dimensional datasets, or datasets that were reduced to two dimensions using Principal Component Analysis (PCA) or some other dimensionality reduction technique. Furthermore, the parameters of this algorithm need to be tested and adjusted for multiple dimensions, to better deal with the curse of dimensionality, but this is a topic to be addressed in the future.

The Gradient-k algorithm is very similar to the original k-means with one main modification and one extra step. The first step of the algorithm is to discretize the point space into evenly spaced boxes along each dimension from the smallest to the largest value. This step is very sensitive to extreme outliers in the data, and as such these points should be removed before this operation. From this point onward, the algorithm will no longer operate in the point space, but in the box space, leading to significant computational advantages when dealing with datasets having a large number of samples. Another advantage of the box space is that, by the nature of the operation, it has the effect of sphering and normalizing the data. Because each dimension is divided into the same number of segments from minimum to maximum, the dimensions are normalized, and the data is stretched and compressed such that the data fits into a square (2D), cube (3D), hypercube (nD). The algorithm then clusters the boxes rather than the points, and then the points are assigned to the clusters to which their box belongs. The second step of the algorithm is to count the number of points in each box and, because each box occupies the same amount of space, this provides a density function of the points.

The third step of the algorithm is to compute the gradient of the density function. Before the computation of the gradient, it is first necessary to apply a smoothing kernel to the density function to remove high spatial frequency noise and keep only the relevant information. The size of this smoothing kernel (*σ*) is one of the tunable parameters and needs to be adjusted for each dataset, such that it does not remove relevant separations between clusters. The result of the first three preprocessing steps is exemplified in Fig 1.

**Fig. 1.**
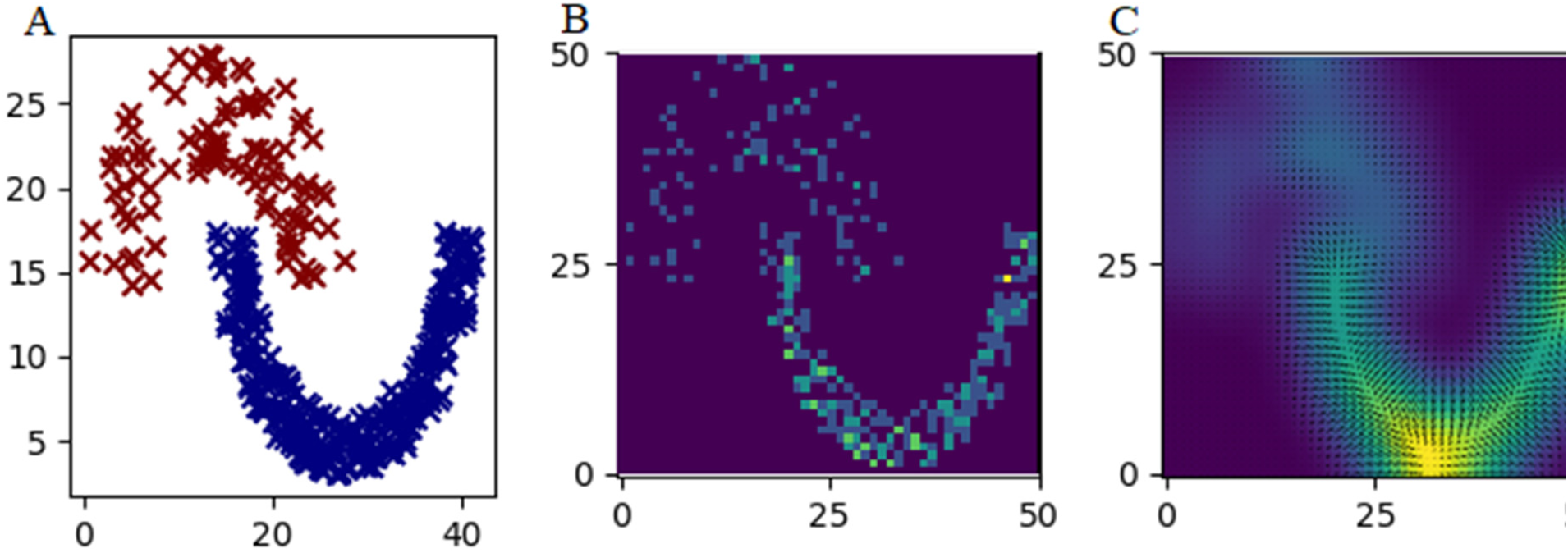
Example of the first two pre-processing steps required for the Gradient-k algorithm. A: the raw points in the data colored using the ground truth labelling. B: the result of the point counting operation for each box. C: the result of the smoothing and gradient computation steps (here the gradients are in their original scale, showing the density difference between the two clusters).

From this step onwards the Gradient-k algorithm operates similarly to k-means, but in box space rather than point space, and with a modification of the distance function. K starting boxes are chosen following the K++ initialization algorithm. The distance from each box to the starting boxes is then computed and multiplied with a correcting factor. The correcting factor is computed by calculating the angle between the gradient of the box and the direction to the cluster centre, dividing the angle with π), and multiplying by an angle importance constant α. A value of 1 is added to the correcting factor, such that the correcting factor is bounded between 1 and α + 1, and the corrected distance is bounded between the original distance and the distance multiplied by α + 1 (Eq. 1, where *θ* is the angle between the direction to the cluster centre and the gradient, *α* is the angle importance constant, *d* is the Euclidean distance, and *d’* is the modified distance).

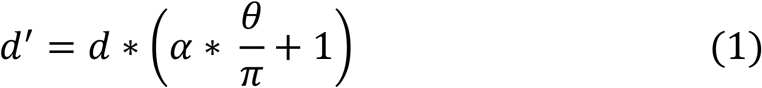

For boxes that are close to a cluster centre, if the direction of density increase leads away from this cluster centre (a large angle between the gradient and direction), the effective distance that the k-means algorithm will consider will be larger, and if the angle is small, the distance will be reduced. The *α* is another tunable parameter and must be adjusted for the dataset. *α* will depend on the number of dimensions that the data has, to correct for the curse of dimensionality. As a function of the resulting effective distances, each box is assigned to the ‘closest’ cluster centre, the cluster centres are recomputed as the weighted average of the coordinates of the boxes in each cluster (here the weights are the number of points in each box), and a new iteration begins. The algorithm stops either when the cluster centres converge (the cluster centres no longer move) or when the cluster centres movement is below a parameter *δ*, which depends on the number of boxes.

Pseudocode for Gradient-k:

~~~
Input: X (m points by n features), N (number of boxes), *α* (angle
importance), *σ* (smoothing kernel standard deviation), *δ* (minimum
cluster displacement), max_epoch (maximum number of iterations)
Pre-processing:
  1. Discretize the space into N boxes along each dimension
  2. Count the number of points in each box, creating a density map
  3. Apply a Gaussian smoothing kernel with standard deviation *σ* on the density map
  4. Compute the gradient of the smooth density space
  5. Choose k initial clusters using K++ initialization and compute coordinates in box space
Main loop (repeat until cluster movement below *δ* or max_epoch iterations are executed:
  6. For each box:
      – compute the distance to each cluster center (d)
      – compute the angle between the density gradient and the direction to each cluster center (*θ*)
      – apply equation 1 to compute the adjusted distance (d’)
      – assign the box to the cluster with the smallest adjusted distance
  7. For each cluster:
      – new centroid = weighted (by number of points in the box) mean
of all boxes assigned to that cluster
~~~

### Parameter Search

For a fair comparison between clustering algorithms, we had to make sure that the algorithm parameters were optimized for the datasets that we were clustering. In the case of Gradient-k, the parameters that we optimized were *α, σ*, and *N*, while in the case of DBSCAN the parameters were *ε* and *minPts*. For this problem, we used the Optuna framework for Python (12), which allows for automatic sampling of a variable space and finding the best parameter combination for a certain problem. We used the TPE (Tree-structured Parzen Estimator) algorithm for sampling the variable spaces, and only defined the range available and the type for each variable (DBSCAN: *ε* as float between 0 and 0.2, *minPts* as integer between 2 and 7; Gradient-k: *N* as integer between 10 and 100, *σ* as float between 0 and 1, *α* as float between 0 and 50).

In the case of DBSCAN, because the algorithm is fully deterministic (has no stochastic elements), one run was evaluated for 2000 parameter combinations or until the algorithm achieved 100% accuracy. In the case of Gradient-k, because the performance can depend on the initialization of the algorithm, we performed 20 runs for each of the 2000 parameter combinations. In order to speed up Gradient-k hyperparameter optimization, we also implemented early stopping: before evaluating the 20 runs with random initializations, one run was evaluated with optimal starting conditions (the initial points were defined as the cluster centers), and the performance was compared to that of other parameter combinations. We only evaluated the 20 runs if the performance with optimal start was greater than or equal to the best performance obtained under optimal starting conditions.

### Comparative Analysis

For the comparative analysis of classic k-means, DBSCAN, and the Gradient-k algorithm, we performed 1000 test runs with the same initialization (for k-means and Gradient-k, respectively). We also performed a set of 1000 tests with the Gradient-k algorithm with an alpha parameter of 0 (the initial points were chosen according to the K++ initialization algorithm, and for the Gradient-k algorithm we chose the boxes in which the points were). The stopping criterion δ was chosen to be 1% of the range of the data for k-means and 1% of *N* for Gradient-k. For each run of the algorithms, we computed the number of iterations until convergence and the accuracy of the resulting clustering.

For benchmarking, we used datasets (13–18) commonly employed to test clustering algorithms. We also used one synthetic dataset (19), generated such that it poses similar issues to one of the most difficult and most often encountered problems in neuroscience: spike sorting (20). Here, the goal is to detect all neuronal spikes in a signal (which can come from one or more implanted electrodes) and assign those spikes to the distinct neurons that generated them. Due to the variable distance from the neurons to the electrode, as well as the neurons producing different waveforms, the waveforms and amplitude are generally used for clustering. In this case, we chose a synthetic monotrode dataset, which had labelled spikes (19). We took the raw data, cut the spikes from the given starting locations (where the spike was inserted into a white noise signal), and aligned them to their maximum amplitude, such that the clustering task was more similar to a real life situation (in most cases, the procedure for spike detection is a statistical thresholding of the data and cutting a window of given length around the peak of the spike).

## Results

As shown Fig. 2, the Gradient-k algorithm outperformed or matched the performance of k-means on all tested datasets. Similarly, when compared to DBSCAN, Gradient-k had better or similar performance on all sets, apart from the Aggregate dataset. In terms of number of iterations, Gradient-k converged in a similar number of iterations to k-means for all datasets except for the spike dataset, where it converged significantly faster. In almost all cases, the differences seen in figure 3 are highly significant (*p* << 0.001, Bonferroni corrected t-tests), except for Gradient-k and k-means differences in: number of iterations on the Unbalanced set (*p* ≈ 0.003) and accuracy on the G2_50 set (*p* ≈ 0.002).

**Fig. 2.**
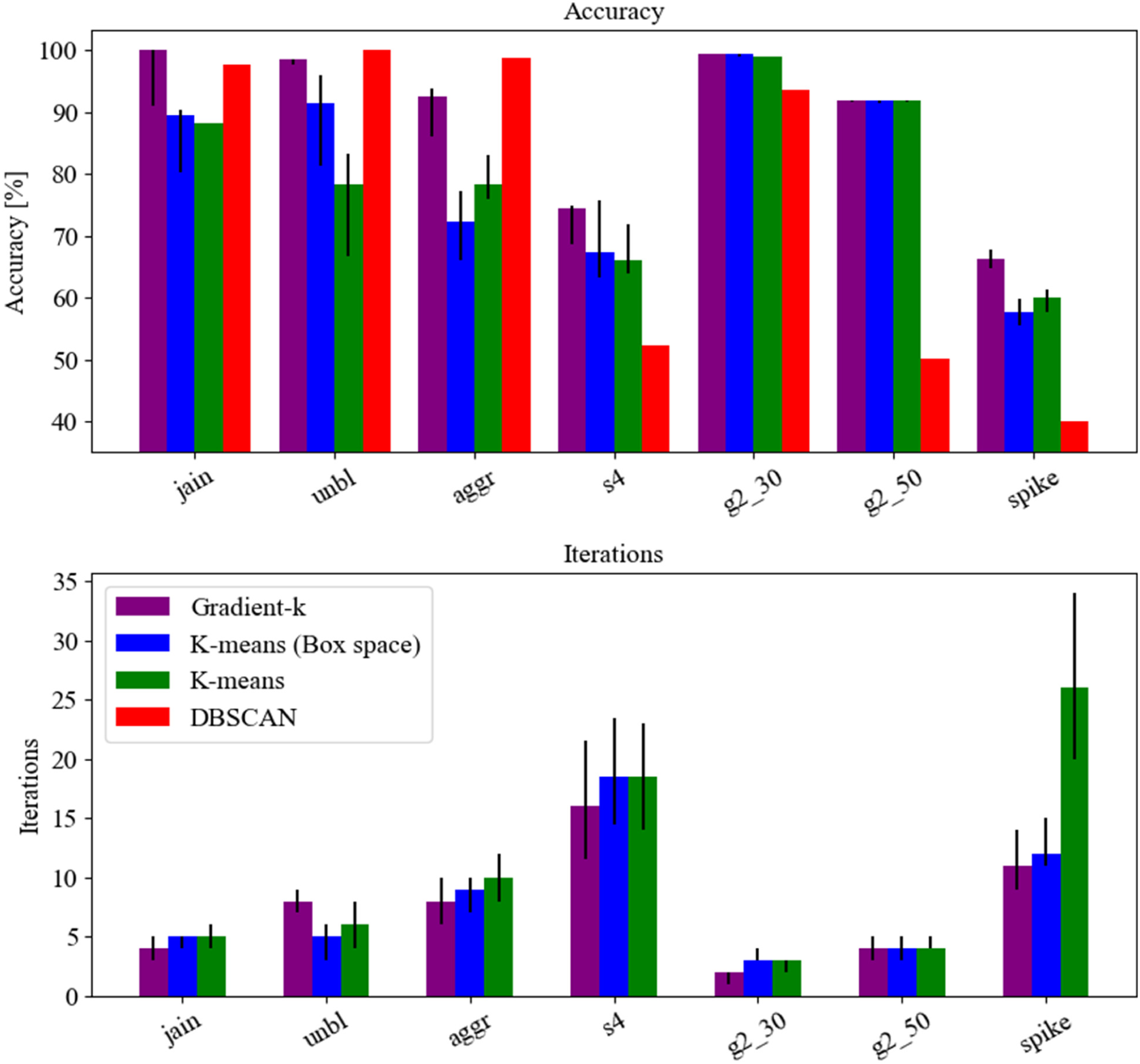
The result of the comparative testing of the algorithms. On the top is the median accuracy expressed in percentages of the algorithms with respect to the true labelling of the datasets. On the bottom, the median number of iterations required for the algorithms to meet the stopping condition. The error bars show Q1 and Q3 quartiles of the data.

**Fig. 3.**
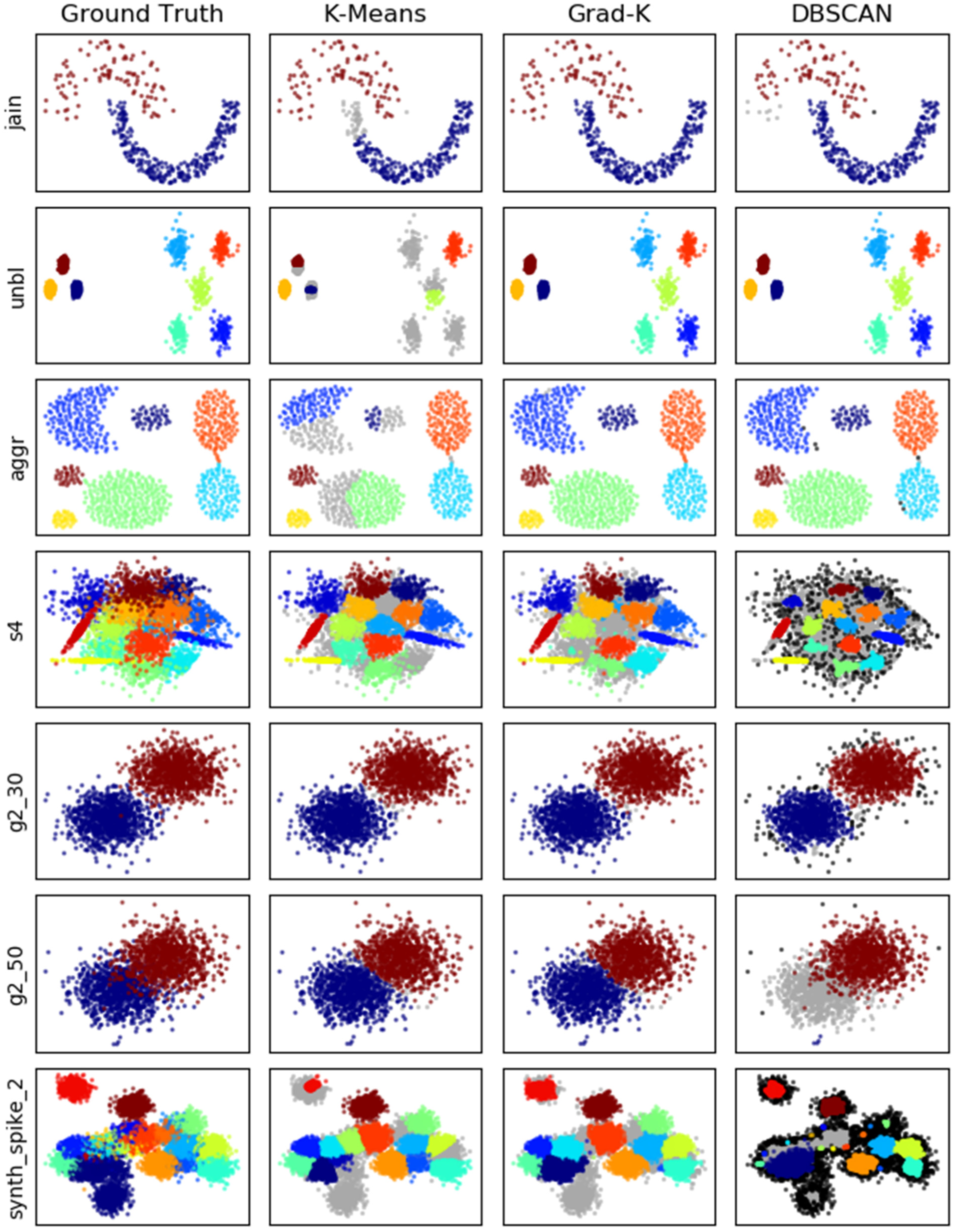
Clustering results from one run of each algorithm, alongside the ground truth for each dataset. Grey points are misclustered and black points are marked as noise by the DBSCAN algorithm.

In figure 3, one can observe that Gradient-k produces clusters that more closely resemble the ground truth than K-means. This is most clearly visible for the datasets where the separation boundary between pairs of clusters cannot be resolved by a straight line (Jain, S4, Synth_spike). In these cases, the additional information provided by the gradient function allowed the algorithm to separate based on density rather than Euclidean distance. Furthermore, this information also allowed the algorithm to delineate between the clusters more accurately in the agglomeration dataset, which have uniform distributions. This is due to the fact that the smooth gradient map that is computed blurs out the edges of the cluster, creating a clear boundary for the algorithm to follow. Finally, the G2-50 set and the Synt_spike set show cases where DBSCAN has worse performance than Gradient-k due to the fact that the inter-cluster space density is too close to the intra-cluster space, leading to the merging of clusters.

Each of the chosen test datasets has specific properties that test the capabilities of the different clustering algorithms. The Jain dataset has a non-linear separation boundary between clusters of different densities, which Gradient-k and DBSCAN can resolve, but k-means cannot. The Unbalance dataset has clusters of high density close together as well as spread out clusters of low density, which leads to the k++ initialization algorithm to choose suboptimal starting points. The Agglomeration dataset poses a problem in which the clusters have uniform (rather than Gaussian) distributions, thus violating a core assumption of the k-means method. The S4 set presents a problem in which Gaussian clusters of different shapes and densities in different directions are close together, thus being a good example of the tessellation problem. Finally, the G2 sets present a problem in which two Gaussian clusters are in two different states of overlap and clearly shows the separation boundary generated by the two algorithms.

## Discussion

### Advantages and disadvantages of Gradient-k

The Gradient-k algorithm has several advantages over existing clustering algorithms and manages to solve many of the problems discussed in this article. First, Gradient-k allows the formation of non-linear splits by clustering the boxes rather than the points and by using the direction of density increase to decide to which cluster the points on the border belong to. Therefore, the algorithm can find clusters with non-gaussian shapes, and the tessellation problem is mitigated. Second, because of the application of a smoothing function over the density space before calculating the gradient of the density function, the algorithm is less sensitive to outliers in between clusters. This smoothing also allows the algorithm to distinguish clusters that are distributed uniformly. The density at the edge of these uniform clusters will be noisy before the smoothing, but the smoothing has the dual effect of minimizing the impact of this noise and having the density function increase from the edge of the cluster towards its centre, informing the algorithm for data at the boundary of the cluster. Third, the proposed correction of the k- means algorithm also has the advantage of allowing it to distinguish clusters of varying densities in the same dataset. The gradients around both dense and less dense clusters are deflected by the greater density of the points in the cluster compared to the inter-cluster space, and because we use only the direction of the density gradient, rather than its magnitude, the Gradient-k algorithm can detect both cluster types at the same time.

Aside from solving these problems, Gradient-k also has some distinct advantages. One of the greatest advantages of the algorithm is that the computational complexity of the clustering procedure does not depend on the number of points being clustered, and as such, the clustering steps take the same amount of time for arbitrarily large datasets. This effect stems from the fact that the clustering is performed through computations in box, rather than point space. While the first part of the algorithm, the discretization of the point space into boxes and computation of the density function, depends on the number of points being counted, the rest of the algorithm operates only on the boxes. Because the number of boxes is fixed, and usually much smaller than the number of points, the total complexity of the algorithm is greatly reduced. A second advantage of using information about the structure of the data while clustering is that in all cases Gradient-k converges in the same or significantly fewer iterations compared to classical k-means. This is likely because, by using the density gradient direction as information in clustering, the cluster centers are pulled naturally to local maxima of the density function, which coincide in most cases with the cluster centers. Indeed, while applying k-means in box space (with *α* set to 0) reduces the number of iterations on its own, we can see that using the angle parameter comes with an extra benefit as well.

In principle, using the gradient of the density function to correct the distance metric could also mitigate the problems that arise in high dimensional spaces. The main issue in high dimensional spaces is that distances carry less information and can no longer reliably be used to measure similarity. On the other hand, the direction of the smoothed density gradient should always be away from inter-cluster space and towards the cluster centre. By making the α parameter in our algorithm larger, the angle between the density gradient and the direction to the cluster centre would eventually matter more than the distance between the point and cluster centre, thus biasing the algorithm in favor of this information.

The algorithm we introduced also has two disadvantages that are shared with the k-means algorithm. First, the number of clusters must be determined *a priori* through different methods, as this is currently a required parameter. This problem could potentially be eliminated by using our smooth density function, but this requires further testing. By counting how many local maxima there are in the smooth density function, it should be possible to determine the number of cluster centres, as well as make a more informed initialization compared with k++. Essentially this could replace the k++ initialization, as well as offer further speed increases by starting with points that are close to the cluster centres. A second drawback is that Gradient-k currently does not scale well computationally to the dimensionality of the dataset. While k-means does not function well in higher dimensions due to the curse of dimensionality, Gradient-k is unfeasible for datasets with high dimensionality because the computational complexity scales exponentially with the dimensionality of the boxes. If we divide each dimension in 100 boxes, then for a 1D problem there are 100 boxes, for a 2D problem there are 100^2^, for a 3D problem 100^3^, and so on.

### Further research and improvements

There are two main avenues for potential improvement of the Gradient-k algorithm. It is necessary to find a way to generalize the algorithm to function in any number of dimensions and to find a rule for how to modify the α parameter as the dimensionality of the problem increases. Furthermore, it is desirable to find a more intelligent initialization, such that the number of cluster centres can be automatically determined, and the location of the starting points can be more informed, in order to increase the speed, accuracy, and to reduce the number of parameters.

The first direction that can be explored is to generalize the algorithm to multiple dimensions and find how the optimum α parameter depends on the dimensionality of the clustering problem. One problem is that the complexity of the algorithm scales exponentially with the dimensionality of the space. It is possible to work around this issue by using PCA to reduce dimensionality, but this does not solve the core problem, because the number of components necessary to fully describe the dataset can still be large enough that the algorithm becomes unfeasible. One potential way that this problem could be mitigated is by ignoring the boxes with few points as being outliers, and automatically eliminating the boxes with no points from the algorithm, essentially reducing the total number of boxes that go into the algorithm. As mentioned earlier, this algorithm could potentially solve the curse of dimensionality, and as such would be desirable to apply in higher dimensions, but we must first reduce the computational complexity for high-dimensional datasets.

The second area that could be improved in Gradient-k is the automatic choice of starting points and number of clusters. In its current implementation, Gradient-k uses the k++ initialization algorithm to choose a desired number (that is set by the user) of initial cluster centres. Manually choosing the number of clusters is not always a straightforward task. In most applications, the number of ‘real’ cluster centres is unknown, and simply looking at the data is not enough. By using the density space to inform this decision, we could eliminate the operator from the equation, make the algorithm easier to use, and have a more informed starting position for the cluster centres, speeding up the algorithm. In k-means clustering, it is possible to ‘warm start’ the algorithm by choosing the starting points manually. In this case, a user would look at the data and guess where the cluster centres are, visually. For some datasets, this is easy, and generally, by doing this k-means converges much faster. If we think about how the users make this decision, it becomes clear that they are essentially starting the algorithm with points situated in high point density locations, and this information can easily be automatically inferred from the density function. Thus, a smart initialization for Gradient-k could eliminate the need to specify the number of clusters and provide a boost to convergence speed similar to how ‘warm starting’ k-means does.

Another option which would allow using the gradient of the density function to correct the Euclidean distance in k-means is kernel density estimation (KDE) (21,22). Similar to the discretization and box counting that we proposed here, KDE allows to infer information about point distribution and density in the space between the given data points. KDE is a procedure through which a sum of Gaussian kernels is computed that are centered in each given point, yielding the resulting density function. Mathematically, the result of this procedure should resemble the output that we get from discretization, box counting, and smoothing, but algorithmically it may be faster and have a lower complexity for high-dimensional data.

On the other hand, a drawback of KDE could be that because it infers the density function without discretizing the space, one would no longer have the advantage of being able to cluster the boxes rather than the datapoints. Furthermore, one would have to use bilinear interpolation (23) to infer the gradient at each datapoint, to assure that the direction of the gradient is appropriate. We would have to test this procedure to see how it compares to the one tested above in terms of accuracy and speed in order to make definitive remarks, but if it outperforms or matches the box counting, it may be a valid alternative for high-dimensional data.

## Conclusions

The Gradient-k algorithm improves the classical k-means algorithm by using additional information in the form of a density function computed from the spatial distribution of the points. The distance that is normally used as a deciding factor in k-means clustering is scaled with a correcting factor based on the angle between the density gradient and the direction towards the cluster centre. This approach allows Gradient-k to avoid many of the pitfalls that the k-means algorithm has, while also offering improvements in terms of speed, due to the nature of the algorithm and the more informed decisions that the algorithm makes. In all tested cases Gradient-k was at least as accurate and at least as fast as k-means. Gradient-k allows non-linear splits, can find clusters of non-Gaussian shapes, and has a reduced tessellation behavior. It can also reliably detect clusters of different densities present in the same dataset and its computational complexity for the clustering step is constant for arbitrarily large datasets (the clustering steps in the algorithm operate in box space). Gradient-k has three main drawbacks in its current form: it still requires *a priori* knowledge of the number of clusters present in the dataset, it requires a parameter search to choose the values for the two tunable parameters, and it scales exponentially with the number of dataset dimensions.

While our current version of Gradient-k has advantages compared to both k- means and DBSCAN, we only tested it on two-dimensional problems, and scaling the algorithm to higher dimensional problems comes with a new set of challenges. Both reducing the computational complexity, and automatically adjusting the α parameter to compensate for the curse of dimensionality would be required to have better performance. Finally, the implementation of an automatic initialization procedure would be highly desirable, both to speed up the algorithm, and to automatically detect cluster candidates, such that the number of clusters is eliminated as a required parameter.

## Acknowledgements

The research leading to these results has received funding from: NO (Norway) Grants 2014-2021, under Project contract number 20/2020 (RO-NO-2019-0504), four grants from the Romanian National Authority for Scientific Research and Innovation, CNCS-UEFISCDI (codes PN-III-P2-2.1-PED-2019-0277, PN-III-P3-3.6-H2020-2020-0109, ERANET-FLAG-ERA-ModelDXConsciousness, and ERANET-NEURON-Unscrambly), and a H2020 grant funded by the European Commission (grant agreement 952096, NEUROTWIN).

## Code availability

Free code implementing Gradient-k is available at: https://github.com/TransylvanianInstituteOfNeuroscience/Gradient_K

## Notes

### Competing Interest Statement

The authors have declared no competing interest.

